# Exposure to graphene oxide sheets alters the expression of reference genes used for real-time RT-qPCR normalization

**DOI:** 10.1101/469304

**Authors:** Irene de Lázaro, Kostas Kostarelos

**Affiliations:** Nanomedicine Lab, Faculty Biology, Medicine and Health and National Graphene Institute, AV Hill Building, The University of Manchester, Manchester M13 9PT, United Kingdom; John A. Paulson School of Engineering and Applied Sciences, Harvard University, Cambridge, MA, 02138, USA; Wyss Institute for Biologically Inspired Engineering, Harvard University, Boston, MA, 02115, USA

**Keywords:** nanomaterials, gene expression, normalization, housekeeping genes

## Abstract

**Background:** studies that unravel the interactions between thin, 2D graphene oxide (GO) sheets and the biological milieu, including cells and tissues, are multiplying quickly as the biomedical applications of those and other 2D materials continue to be explored. Many of such studies rely on real-time RT-qPCR as a powerful, yet relatively simple technique to determine gene expression. However, a systematic investigation of potential GO-induced changes in the expression of reference genes, crucial for appropriate normalization of qPCR data that ensures reliability of the results, is still lacking. In this study, we aimed to cover this gap by investigating the stability of the expression of ten (10) candidate reference genes upon exposure to increasing, but subtoxic, concentrations of GO, with two established algorithms (Bestkeeper and NormFinder). The study was performed in a human cancer cell line (MCF7) and in mouse, non-cancerous primary cells (mouse embryonic fibroblasts, MEFs), to assess different behaviors between cell types.

**Results:** Bestkeeper and NormFinder algorithms evidenced significant deviations in the expression of various reference genes. Ribosomal proteins scored among the most significantly dysregulated targets in both cell types. Expression of *ACTB* and *GAPDH*, the most frequent calibrators in real-time RT-qPCR studies, was also affected, although differences existed between cell lines.

**Conclusions:** this study illustrates the need to validate reference genes for appropriate real-time RT-qPCR normalization, according to specific experimental conditions, when GO-cell interactions occur.

## Background

The use of two-dimensional (2D) nanomaterials in biomedical research is escalating at a fast pace, thanks to their long list of outstanding physicochemical properties (1). Among them, graphene oxide (GO), the oxidized version of graphene, is the candidate of choice in studies that explore direct interactions with the physiological milieu or at the intracellular level. This preference is due to the presence of multiple oxygen functionalities that grant the material stability in aqueous dispersion, and thus, in most biological fluids (2). The abundance of functional groups available for further functionalization in the characteristically large surface area of GO (larger than that of other nanomaterials due to the bi-planar structure) also turns these flat flakes into versatile platforms to accommodate, transport and deliver a wealth of molecules of biological relevance (3).

However, the effects that may be triggered in the physiology and molecular machinery the cell upon contact with GO are not yet fully understood. Although its cytotoxicity profile is thought to be privileged compared to that of other nanoscale materials — at least that of small GO flakes, of lateral dimensions < 1 μm, produced under highly controlled conditions — GO is certainly known not to be inert to cells and tissues (4). Changes in gene and protein expression have been reported that depend strongly on the physicochemical characteristics and dimensions of the material but also on the cell type or tissues under investigation and overall conditions surrounding the material-cell interaction (5-8).

Real-time, reverse transcription, quantitative polymerase chain reaction (RT-qPCR) is a powerful technique to determine the expression levels of a specific mRNA. Its ubiquitous use is justified by its high speed, sensitivity, specificity and reproducibility; but also by its relative simplicity compared to the technical, analytical and economical demands of deep sequencing (9). However, correct normalization of relative data is critical for the reliability of the technique and has been the topic of numerous studies covering both mathematical approximations and the adequacy of calibrators (10-12).

To account for possible differences in the amount of starting material, RNA recovery and integrity, efficiency of cDNA synthesis and overall transcriptional activity of the cell or tissue of interest, all of which can easily compromise the accuracy of the analysis if not properly corrected, so-called “reference” or “housekeeping genes” are used as calibrators to normalize real-time RT-qPCR data. Traditionally, those are selected among genes involved in very fundamental cellular functions (e.g. transcription, translation, cytoskeletal structure and other basic metabolic pathways) under the assumption that they are constitutively expressed across different cell types, tissues and conditions (13). However, several reports have unveiled the lack of stable expression of many of such traditionally considered housekeeping genes under a number of circumstances, including different cell phenotypes and tissues (14, 15), physiological and diseased states (16-18) and experimental conditions (19, 20). MIQE guidelines, released in 2009 to promote transparency and good practice in RT-qPCR studies, strongly advise against the standardization of gene expression data with a single, non-validated, reference gene (21). Several algorithms and software tools, including Bestkeeper (22), NormFinder (23) and GeNorm (24), have been developed to facilitate the validation of intended calibrators.

It is of special concern that the impact of GO exposure in the expression of commonly used reference genes remains, to our knowledge, unexplored. Even more alarming is the absence of such validation in a number of studies, reviewed elsewhere (25), where GO was used as a component of siRNA and mRNA delivery vectors and that relied on RT-qPCR to evaluate the efficacy of the silencing or forced gene expression achieved.

In this study, we aimed to assess the stability of the expression of ten candidate reference genes (**Tables 1 and 3**) upon *in vitro* exposure to sub-cytotoxic concentrations of highly characterized, endotoxin-free, GO flakes. We performed the study in the human cancer cell line MCF7 and in murine, non-cancerous, primary cells (mouse embryonic fibroblasts, MEFs), to investigate if the impact of GO in gene expression depended on cell type. Real-time RT-qPCR analysis in this work was performed in strict compliance with MIQE guidelines to ensure the reliability of the results, and Bestkeeper and NormFinder algorithms were used to rank the performance of the candidates as stable reference genes. A more in-depth analysis of the impact of GO exposure was performed for the most dysregulated genes in each cell type.

## Results

### Selection of candidate reference genes

We based our selection of candidate reference genes on a literature search to identify those most commonly used in real-time RT-qPCR normalization and on a meta-analysis conducted by de Jonge et al (26) that analyzed 13,629 human and 2,543 mouse gene arrays from previous publications. De Jonge et al identified various novel housekeeping genes — including *RPS13*, *RPL27*, *RPL30* and *OAZ1* — whose expression proved significantly more stable than that of more commonly used calibrators, such as *GAPDH* and *ACTB.* We paid special attention to select candidates with different cellular functions, to minimize bias introduced by co-regulated genes. In agreement, ribosomal proteins (*RPS13*, *RPL30* and *RPL27*), metabolic enzymes (*OAZ1*, *GAPDH*, *UBC*, *HMBS*), transcription factors (*TBP*), kinases involved in signaling pathways (*MAPK1*) and structural genes (*ACTB*) were all included (**Tables 1 and 3**).

### RNA samples and RT-qPCR reactions

To study the impact of the material on the expression of candidate reference genes, MCF7 cells and MEFs were exposed to increasing concentrations of endotoxin-free GO (0, 5, 10 and 50 μg/ml). This range was selected based on previous studies from our laboratory that confirmed lack of toxicity in a number of cell lines (data not shown). Full characterization of the material, produced *in house* by a modified Hummer’s method as previously described (27, 28), has been reported in a previous publication (29) and the most relevant parameters are summarized in **Table S1**. In brief, lateral dimensions did not surpass the 2 μm threshold and thickness corresponded to 1-2 single GO layers. Functionalization degree was estimated as 41%. Exposure took place in the absence of FBS for the first 4 h, to mimic conditions commonly used when testing nanomaterials *in vitro.* Gene expression was assessed 24 h after the initial exposure by real-time RT-qPCR. All procedures were performed in strict compliance with MIQE guidelines (21), to ensure reliability of the results. To provide the transparency required by these recommendations, a MIQE guidelines checklist is provided in **Table S2**. To avoid error introduced by poor RNA quality, A_260/A280_ and A_260/230_ ratios of all RNA samples included in the study ranged between 1.70 and 2.1, their RIN values were > 8.9, and their 28S/18S ratios ranged between 2.0-3.9 (**Tables S3 and S4**). Primers, designed *in house* for the study, amplified all transcription variants of each target with equal product length. Their details are given in **Tables S5 and S6**, including the efficiencies of qPCR reactions (E), determined by serial dilution of the cDNA template.

### Expression stability of candidate reference genes in MCF7 cells exposed to GO

We first used Bestkeeper software (22) to obtain preliminary information regarding the expression of each candidate reference gene in MCF7 cells treated with GO. Bestkeeper provides the standard deviation (SD) and the coefficient of variance (CV) of the quantification cycles (Cq, expressed as crossing point (CP) in the original software), for each gene. CV is calculated as the percentage of the Cq SD to the Cq mean. Genes with SD>1 are considered to have an unacceptable range of variation (22). As reported in **Table 2**, all gene candidates showed Cq SD<1, thus none had to be excluded from further analysis on such grounds. However, *β*-*Actin* (*ACTB*) clearly scored as the less stably expressed gene among all candidates, with SD=0.58 and CV=4.15 (**Table 2**). It was followed by all three ribosomal proteins investigated: *RPL27* (SD=0.41, CV=2.37), *RPS13* (SD=0.25, CV=1.42) and *RPL30* (SD=0.30, CV=1.39). The enzyme Ornithine decarboxylase antizyme 1 (*OAZ1*) showed also a relatively high variability of expression (SD=0.27, CV=1.34), compared to other candidates. *GAPDH*, *MAPK1*, *UBC* and *HMBS* were ranked as the most stable genes, based on lower SD and CV values (see **Table 2**).

**Table 1.** Candidate reference genes included in MCF7 study (human).

| Gene symbol | Gene name | Gene ID | Protein function |
| --- | --- | --- | --- |
| ***RPS13*** | Ribosomal protein S13 | 6207 | Translation |
| ***RPL27*** | Ribosomal protein L27 | 6155 | Translation |
| ***RPL30*** | Ribosomal protein L30 | 852853 | Translation |
| ***OAZ1*** | Ornithine decarboxylase antizyme 1 | 4946 | Metabolic enzyme |
| ***ACTB*** | β-actin | 60 | Structural |
| ***GAPDH*** | Glyceraldehyde-3 phosphate dehydrogenase | 2597 | Metabolic enzyme |
| ***MAPK1*** | mitogen-activated protein kinase 1 | 5594 | Signalling |
| ***UBC*** | Ubiquitin C | 7316 | Metabolic enzyme |
| ***HMBS*** | Hydroxymethylbilane synthase | 3145 | Metabolic enzyme |
| ***TBP*** | TATA-box binding protein | 6908 | Transcription |

**Table 2.** Descriptive statistics from Cq values extracted from Bestkeeper algorithm, MCF7
study. Geometric mean (geoMean), standard deviation (SD) and coefficient of variance (CV) of Cq data from ten candidate reference genes in MCF7 cells exposed to increasing concentrations of GO (n=12). CV is calculated as the percentage of the Cq SD to the mean Cq.

| Gene | geoMean [Cq] | SD [±Cq] | CV [%Cq] |
| --- | --- | --- | --- |
| ***RPS13*** | 17.70 | 0.25 | 1.42 |
| ***RPL27*** | 17.51 | 0.41 | 2.37 |
| ***RPL30*** | 21.66 | 0.30 | 1.39 |
| ***OAZ1*** | 20.04 | 0.27 | 1.34 |
| ***ACTB*** | 14.02 | 0.58 | 4.15 |
| ***GAPDH*** | 15.48 | 0.17 | 1.07 |
| ***MAPK1*** | 20.07 | 0.19 | 0.95 |
| ***UBC*** | 24.09 | 0.18 | 0.75 |
| ***HMBS*** | 19.33 | 0.19 | 1.00 |
| ***TBP*** | 21.28 | 0.22 | 1.05 |

Bestkeeper assesses stability of expression based on the two parameters above (SD and CV), but does not offer a strict ranking of the most stable housekeeping genes. It provides a normalization index (Bestkeeper index) to normalize each sample, calculated as the geometric mean of Cqs from the best performing candidates (22). We used the model-based NormFinder approach (23) to obtain a precise ranking. In this case, the algorithm accounts for all different experimental groups included in the study and considers intra- and inter-group variation to provide a direct measure of stability, defined as *stability value.* The lower this value, the higher the stability of expression of the candidate (23). In agreement with Bestkeeper data, *ACTB* and the three ribosomal proteins (*RP27*, *RPL30* and *RPS13*) scored the highest (less stable) values (0.367, 0.264, 0.230 and 0.136, respectively) (**Figure 1**). *GAPDH* was ranked as the most stably expressed gene (stability value=0.029). NormFinder also defines the stability value for the best combination of two reference genes. In this case, the combination of *GAPDH* and *HMBS* did not improve, but obtained the same stability value as *GAPDH* alone, 0.029 (**Figure 1**).

**Figure 1.**
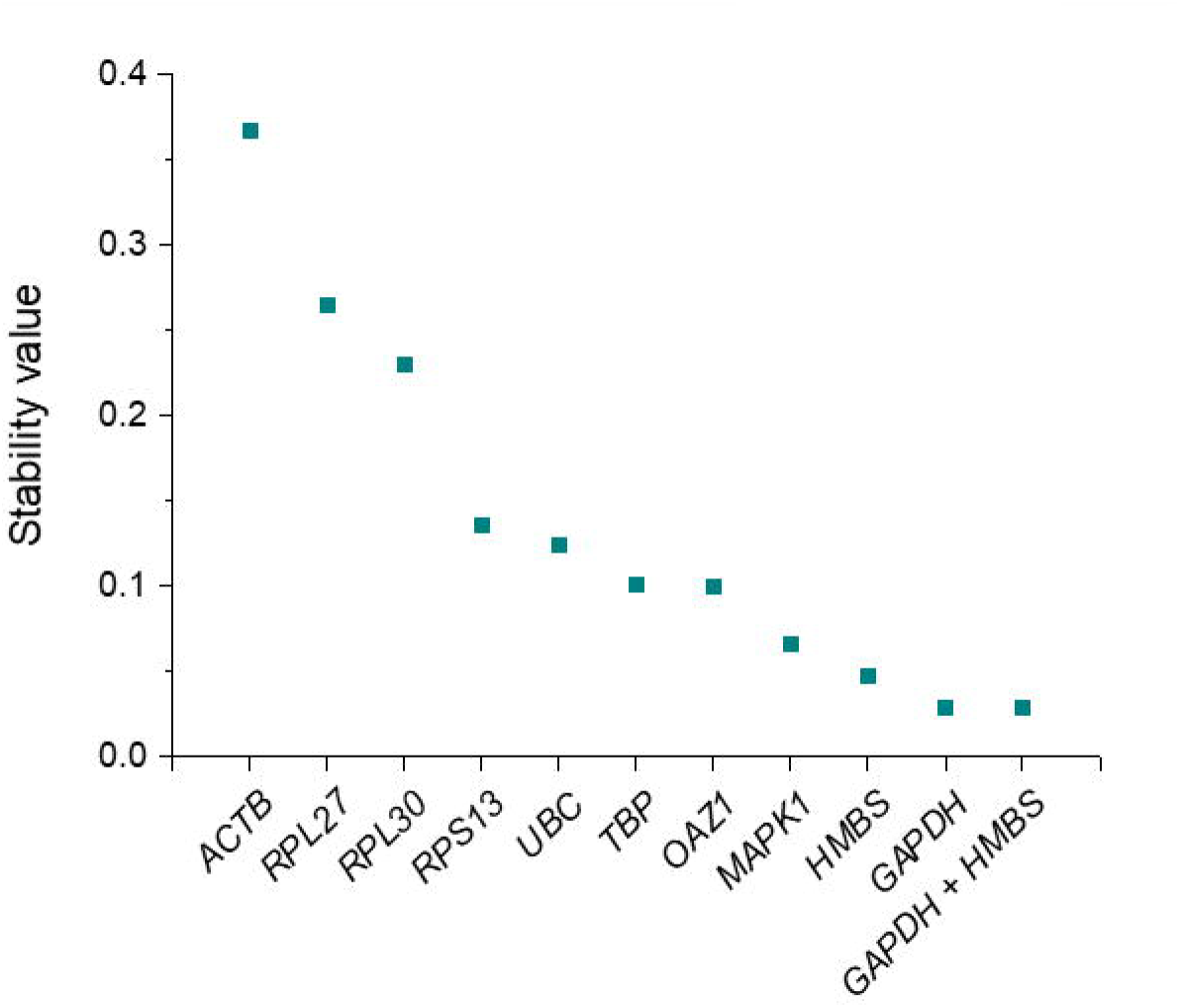
Stability values of ten reference genes in MCF7 cells exposed to GO, calculated with NormFinder algorithm. Candidate reference genes are ranked according to decreasing stability values (i.e. increasing stability of expression). *GAPDH* was ranked as the most stable gene. The most stable combination of two genes (*GADPH* + *HMBS*) is also represented. See **Table 1** for gene names.

Overall, these data highlighted differences in the stability of expression among various common reference genes utilized for RT-qPCR data normalization, when MCF7 cells were treated with different concentrations of GO.

### Expression stability of candidate reference genes in MEFs exposed to GO

To confirm whether the observations above where exclusive to MCF7, a human cancer cell line, we repeated the study with non-cancerous, primary mouse cells. MEFs were isolated from E12.5 embryos from the CD1 background, cultured for no more than three passages, and treated under the same conditions used in the MCF7 study. The homologue genes were evaluated as candidate references, since they are conserved in the mouse and human genomes (**Table 3**), but specific primer pairs were designed (Table S6). Descriptive statistics from Cq values retrieved from Bestkeeper software are summarized in **Table 4**. The mean Cq values of several candidates differed from those registered in MCF7 cells (**Table 2**), which underlines the need to validate the calibrator of choice when comparing gene expression in different cell types. As in the MCF7 cell line, *Actb* Cqs showed the highest variation among all candidates in MEF samples (SD=0.21, CV=1.47). However, SD and CV values were generally lower that those observed in MCF7 cells, which evidenced an overall higher stability of expression of the candidates upon GO treatment. The same observation was confirmed by NormFinder analysis, whereby the stability values of all candidates ranged between 0.172 and 0.071 in MEFs (**Figure 2**), while they spanned from 0.367 to 0.029 in the MCF7 study (**Figure 1**). This algorithm identified *Rpl27* as the most unstable gene in MEFs exposed to GO, with a stability value of 0.172, and closely followed by *Gapdh* (0.154). On the opposite side of the ranking, *Hmbs* was the most stable candidate (stability value=0.071), but the combination of *Rpl30* and *Tbp* provided an even lower stability value (0.041) (**Figure 2**).

**Table 3.** Candidate reference genes included in MEF study (mouse).

| Gene symbol | Gene name | Gene ID | Protein function |
| --- | --- | --- | --- |
| ***Rps13*** | Ribosomal protein S13 | 68052 | Translation |
| ***Rpl27*** | Ribosomal protein L27 | 19942 | Translation |
| ***Rpl30*** | Ribosomal protein L30 | 19946 | Translation |
| ***Oaz1*** | Ornithine decarboxylase antizyme 1 | 18245 | Metabolic enzyme |
| ***Actb*** | β-actin | 11461 | Structural |
| ***Gapdh*** | Glyceraldehyde-3 phosphate dehydrogenase | 14433 | Metabolic enzyme |
| ***Mapk1*** | Mitogen-activated protein kinase 1 | 26413 | Signalling |
| ***Ubc*** | Ubiquitin C | 22190 | Metabolic enzyme |
| ***Hmbs*** | Hydroxymethylbilane synthase | 15288 | Metabolic enzyme |
| ***Tbp*** | TATA-box binding protein | 21374 | Transcription |

**Table 4.** Descriptive statistics from Cq values extracted from Bestkeeper algorithm, MEF study. Geometric mean (geoMean), standard deviation (SD) and coefficient of variance (CV) of Cq data from ten candidate reference genes in MEFs exposed to increasing concentrations of GO (n=12). CV is calculated as the percentage of the Cq SD to the mean Cq.

| Gene | geoMean [Cq] | SD [±Cq] | CV [%Cq] |
| --- | --- | --- | --- |
| ***Rps13*** | 16.46 | 0.16 | 1.00 |
| ***Rpl27*** | 27.56 | 0.28 | 1.01 |
| ***Rpl30*** | 21.84 | 0.21 | 0.95 |
| ***Oaz1*** | 17.38 | 0.17 | 0.95 |
| ***Actb*** | 14.16 | 0.21 | 1.47 |
| ***Gapdh*** | 15.44 | 0.17 | 1.10 |
| ***Mapk1*** | 19.56 | 0.21 | 1.05 |
| ***Ubc*** | 20.33 | 0.12 | 0.61 |
| ***Hmbs*** | 21.20 | 0.17 | 0.82 |
| ***Tbp*** | 21.93 | 0.16 | 0.72 |

**Figure 2.**
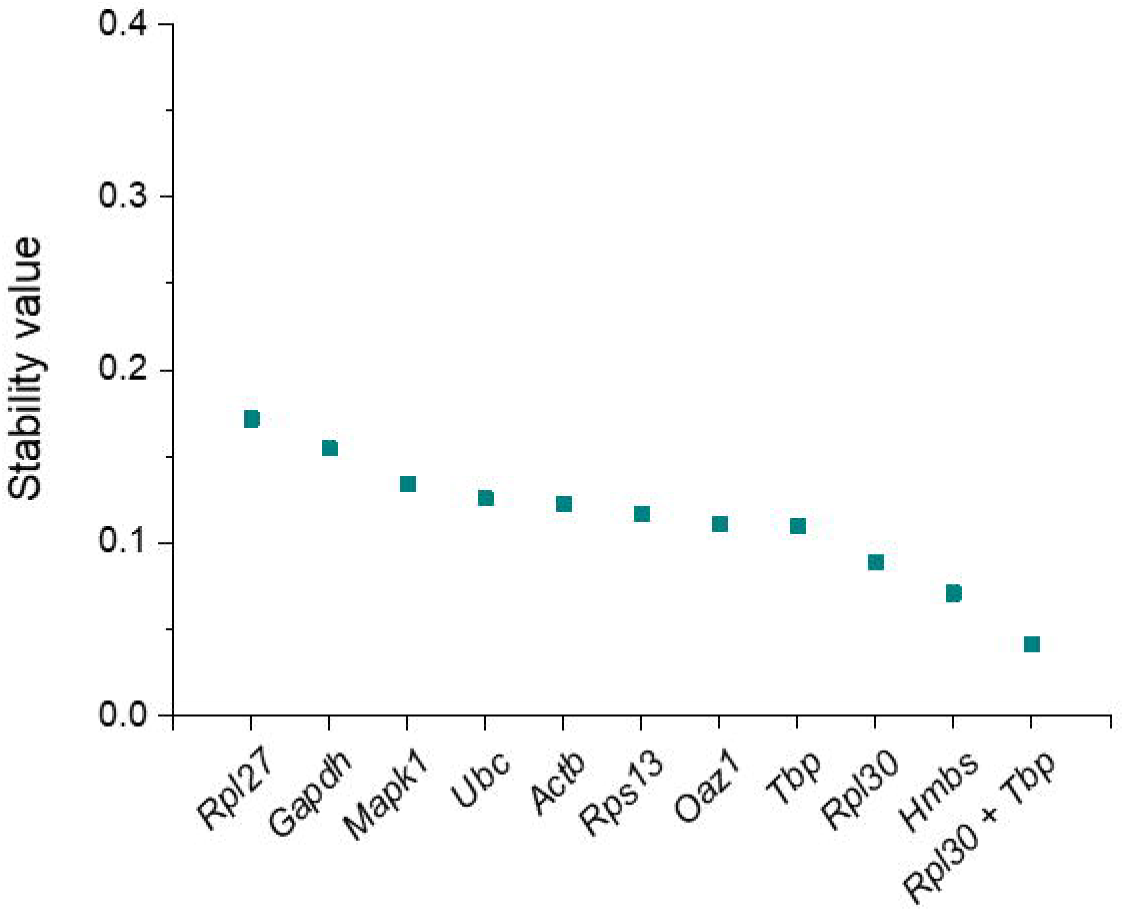
Stability values of ten reference genes in MCF7 cells exposed to GO, calculated with NormFinder algorithm. Candidate reference genes are ranked according to decreasing stability values (i.e. increasing stability of expression). *Hmbs* was ranked the most stably expressed single gene, but the combination of (*Rpl30* + *Tbp*) provided an even lower stability value. See **Table 3** for gene names.

### GO treatment induces dose-dependent *RPL27* downregulation in both cell types

Moved by the poor stability scores of *RPL27* in both MCF7 cells and MEFs, we decided to investigate more closely the relationship between the GO treatment and the mRNA levels of this gene. In MCF7 cells, *RPL27* expression was normalized to the geometric mean of *GAPDH* and *HMBS* Cqs, the best combination of two reference genes inferred from NormFinder analysis (**Figure 3, a**). Following the same algorithm, *Rpl30* and *Tbp* were used as calibrators to normalize *Rpl27* data in MEFs (**Figure 3, b**). In both cases, we found a dose-dependent and statistically significant downregulation of the target in the presence of GO, which confirmed the inadequacy of *RPL27* as reference gene under such experimental conditions. The dysregulation was more pronounced in MCF7 cells, where exposure to the lowest concentration of GO tested (5 μg/ml) already induced an 18% downregulation of *RPL27* (p=0.03). At 50 μg/ml, the downregulation reached 56% (p=0.00005). In MEFs, however, the latter concentration was the only one to produce a statistically significant change in *Rpl27* expression (25% downregulation, p=0.02). Data normalization using the Bestkeeper index confirmed these results (**Figure S2**).

**Figure 3.**
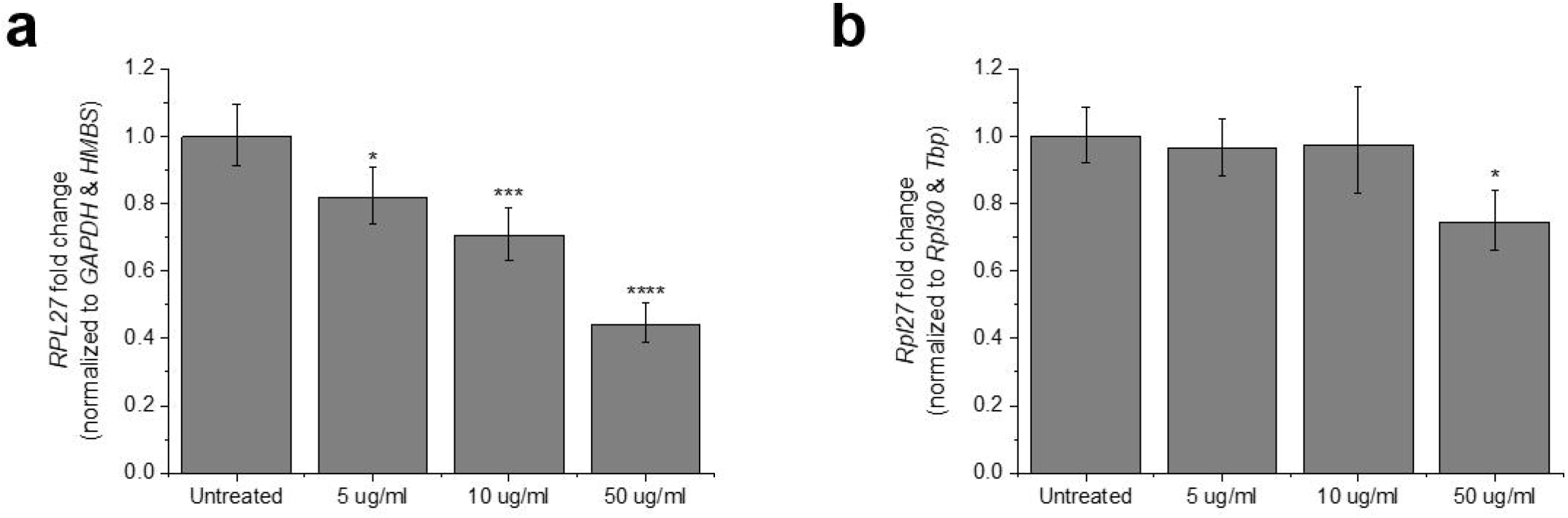
Dose-dependent downregulation of *RPL27* in MCF7 cells and MEFs exposed to GO. **(a)** Relative *RPL27* expression normalized to the geometric mean of *GAPDH* and *HMBS* Cqs, according to NormFinder ranking of stability values. **(b)** Relative *Rpl27* expression normalized to the geometric mean of *Rpl30* and *Tbp* Cqs, according to NormFirnder ranking of stability values. Bars represent fold change; error bars represent propagation of standard error (SE). ^∗^p<0.05, ^∗∗∗^p<0.001, and ^∗∗∗∗^<p<0.0001, assessed by one-way ANOVA and Tukey’s test, n=3.

### Archetypal housekeeping genes are dysregulated in the presence of GO

*ACTB* and *GAPDH* are without doubts the two genes most frequently utilized to normalize real-time RT-qPCR data (13). However, various studies have called to question their status as housekeeping genes, based on significant fluctuation of their expression under specific treatment or experimental conditions and even among different tissues of the same organism (16, 30, 31). Indeed, in de Jonge et al’s ranking of stable human reference genes, *ACTB* and *GAPDH* occupied positions 57 and 139, respectively (26). In our study, *ACTB* was ranked the most unstable candidate reference gene in MCF7 cells treated with GO, by both Bestkeeper and NormFinder algorithms (**Figure 1, Table 2**). In this cell line, *ACTB* mRNA levels were significantly upregulated in the presence of 50 pμ/ml GO (p=0.04, **Figure 4a**). In MEFs, *Actb* showed the highest Cq CV (**Table 4**), albeit this variation did not translate into significant changes in the expression of the transcript (**Figure 4c**). *GAPDH* returned even more discrepant results between the two cell types. It proved the most stable candidate in MCF7 cells, according to NormFinder results (**Figure 1**), and consequently its mRNA relative levels remained stable across all experimental groups (**Figure 4b**). However, the same gene scored the second highest stability value (0.154) upon NormFinder analysis (**Figure 2**) and showed the second highest Cq CV in Bestkeeper (**Table 4**). Indeed, 50 μg/ml GO induced a 1.44-fold *Gapdh* upregulation (p=0.001, **Figure 4d**). All such results were confirmed when Bestkeeper index was used as normalization factor (**Figure S3**). Overall, these findings corroborate the inadequacy of *ACTB* and *GAPDH* as reference genes under particular experimental conditions in which GO is involved.

**Figure 4.**
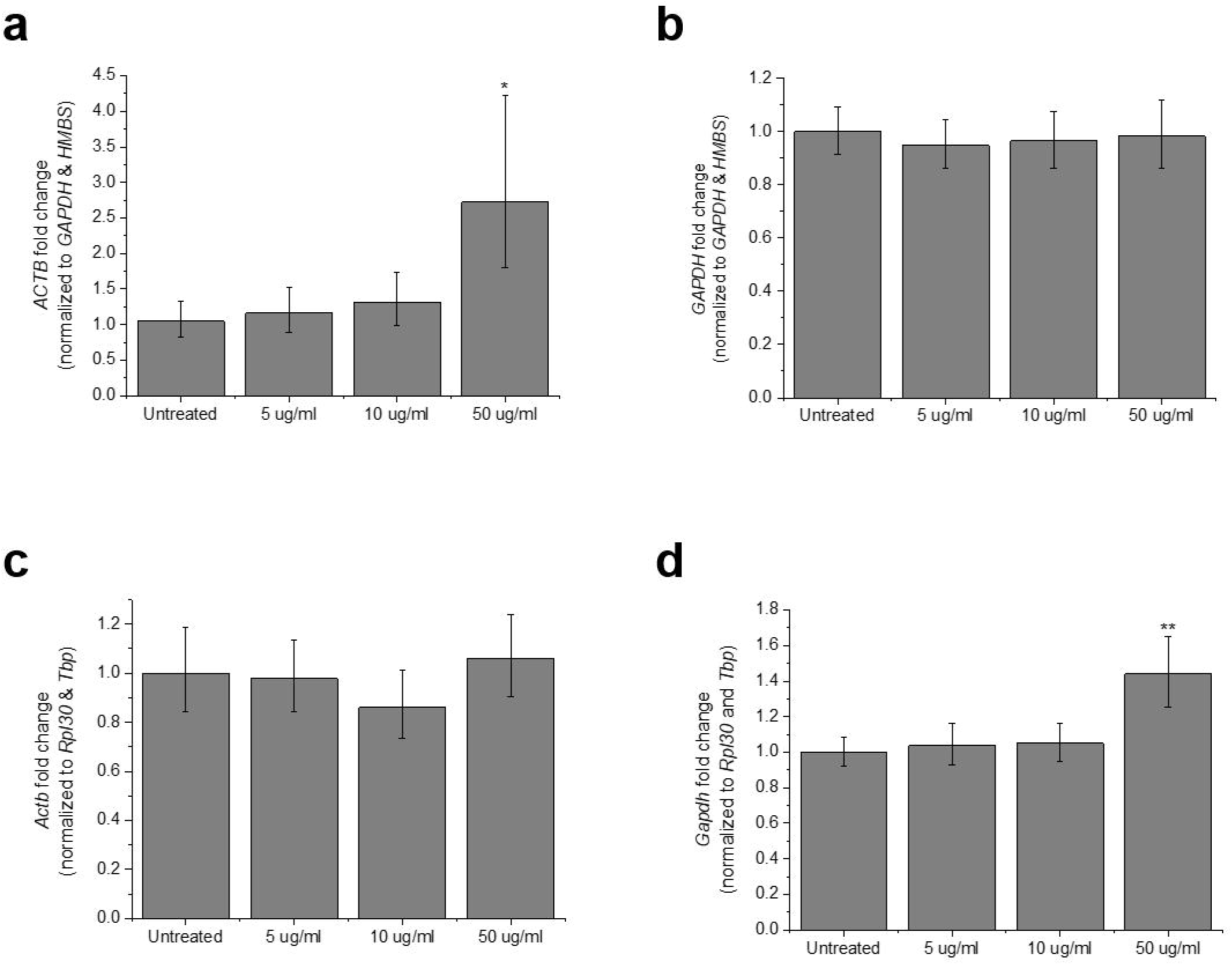
Relative expression of archetypic housekeeping genes in MCF7 cells and MEFs exposed to GO. In MCF7 cells, **(a)** *ACTB* and **(b)** *GAPDH* expression were normalized to the geometric mean of *GAPDH* and *HMBS* Cqs. In MEFs, **(c)** *Actb* and **(d)** *Gapdh* mRNA levels were normalized to the geometric mean of *Rpl30* and *Tbp.* The combination of two reference genes was selected according to NormFinder results. Bars represent fold change; error bars represent propagation of standard error (SE). ^∗^p<0.05 and ^∗∗^p<0.01, assessed by one-way ANOVA and Tukey’s test, n=3.

## Discussion

We have shown here that GO induces significant changes in the expression of common reference genes, at the mRNA level. This is, to our knowledge, the first study to systematically explore the impact of GO exposure in the stability of traditionally considered housekeeping genes, a gap that needed to be filled considering the ever-growing number of studies exploring the GO-cell biology interface. Strikingly, the validation of suitable reference genes was so far absent even in studies in which real-time RT-qPCR data constitutes a central readout of the GO-cell interaction. Most of such works relied on *GAPDH*, exclusively, as calibrator, without data that supported the stability of its expression under their specific experimental conditions (32-36). Another study did not even specify the reference used for normalization, nor whether it had been validated (37).

The above is a matter of concern because (a) MIQE guidelines strongly discourage the use of a single, non-validated gene for standardization (21) and (b) the inadequacy of *GADPH* as calibrator has been evidenced by a number of studies that detected large variations of its mRNA levels across tissue types and under different experimental conditions (16, 30). Our results indeed support these observations and the current thought that a specific gene may behave as stable reference in some, but not all, scenarios. *Gapdh* mRNA levels were significantly upregulated by high (but subtoxic) concentrations of GO in MEFs (**Figure 4d**, **Figure S3d**), rendering it of no use as calibrator. However, this was not the case in MCF7 cells, in which *GAPDH* scored as the most stable candidate reference gene (**Figure 1**). A similar scenario was found for *ACTB*, also a very popular and widespread calibrator that has received increased attention due to the inconsistency of its expression (16). *ACTB* showed the highest stability value, according to NormFinder analysis, in MCF7 cells (**Figure 2**) and its expression significantly increased with GO exposure (**Figure 4a**, **Figure S3a**).

One can also infer from our results that ribosomal proteins are, overall, not appropriate calibrators to normalize real-time RT-qPCR data in the presence of GO. All such candidates included in our study, *RPL27*, *RPL30* and *RPS13*, scored the highest instability in MCF7 cells, in that order and only surpassed by *ACTB* (**Figure 1, Table 2**). *RPL27*, in particular, experienced significant downregulation in the presence of GO, which was dose-dependent and confirmed in both cell types, MCF7 and MEFs (**Figure 3**). These results oppose those reported in de Jonge et al’s meta-analysis, where *RPS13*, *RPL27* and *RPL30* occupied top positions (first, second and fourth, respectively) in the stability ranking of candidate housekeeping genes (26). This discrepancy highlights even more the necessity to validate the reference genes of choice in the presence of GO, as interaction with the nanomaterial seems to dysregulate genes that are otherwise rather stably expressed across an ample variety of conditions.

While *RPL27* mRNA levels followed the same trend in MCF7 cells and MEFs exposed to GO, we also found important differences in the impact of the material in the gene expression profiles of both cell types. First, instability introduced in the expression of candidate reference genes was more pronounced in MCF7 cells compared to MEFs, in which Cq variation was narrower (see SD and CV values in **Tables 2 and 4**, and stability values in **Figures 1 and 2**). Second, we found differences in the ranking of most stable housekeeping genes – even when the same algorithm was used – with stable references in one cell type scoring very poorly in the other (see for instance *GAPDH* data in **Figures 1 and 2**, as discussed above). The discrepancy between cell lines stresses the absolute requirement to validate housekeeping genes under specific experimental conditions. Indeed, inconsistencies in the expression of traditionally considered reference genes between different cell types have already been described, even when such cells were not subjected to any exogenous treatment (14, 15). In addition, GO, as well as other nanomaterials, is known to interact differently with different cell types, including in what concerns to gene expression (7). Given the many parameters that can alter gene expression, our study does not intend to provide a list of reference genes to be used in *in vitro* GO studies, but to highlight the necessity of their systematic validation under specific experimental conditions.

We also found differences within the rankings provided by Bestkeeper and NormFinder (**Figures 1 and 2, Tables 2 and 4**). Such discrepancies were expected, based on the different algorithms that support each software, and have indeed been reported by other studies prior to ours (14, 15, 18, 38). However, we show here that choosing NormFinder or Bestkeeper index for normalization did not change the results obtained for the relative expression of various targets (**Figures 3, 4, S2 and S3**), which grants additional significance to the results shown here.

In spite of the GO-induced changes in gene expression shown here, it is nevertheless to be said that the instability inferred by the material was less pronounced than that triggered by other physiological or experimental conditions. In our study, the highest stability value recorded was 0.367 for *ACTB* in MCF7 cells and 0.172 for *Rpl27* in MEFs (**Figures 1 and 2**), in the range of the values that Lemma et al observed among cancer stem cells (15). However, Ali et al reported stability values above 0.6 when comparing different human lung cancer cell lines (14) and the same maximum value was observed by Wierschke et al in a study that compared brain tissue samples in epileptic patients and healthy controls (18). This comparison, however, does not eliminate the need for appropriate validation of reference genes, since the GO-induced changes in their expression that we have reported here would introduce statistically significant errors in the normalization of relative gene expression data.

## Conclusion

We have shown here that *in vitro* exposure to GO sheets alters the expression of various candidate reference genes used to normalize real-time RT-qPCR data, including very frequently used calibrators such as *GAPDH* and *ACTB.* We have also demonstrated that the magnitude and nature of the changes induced vary between different cell types and therefore reference gene validation cannot be extrapolated but must be specifically determined according to defined experimental conditions. Those may include, but may not be limited to, physicochemical characteristics of the material, dose, exposure conditions and cell type of interest. Using stable reference genes is imperative to obtain reliable gene expression data.

## Methods

### Graphene oxide

GO was produced *in house* following a modified Hummer’s method as previously described(27, 28). Full characterization of this material was reported in a previous publication(29), where it was termed *small* GO (s-GO) in order to differentiate it from larger GO flakes that have not been used in the present study. A summary of this information is provided in **Table S1**. In brief, the lateral dimensions of GO flakes do not surpass 2 μm and their thickness corresponds to 1-2 layers of GO (1-2 nm). The degree of functionalization was estimated as 41% by thermogravimetric analysis (TGA). Presence of oxygenated functionalities in form of hydroxyls, carboxyls and epoxides was confirmed via x-ray photoelectron spectroscopy (XPS). Surface charge was strongly negative (ζ=-55.9 ± 1.4 mV).

### Primary cell extraction, cell lines and culture

The MCF7 human breast cancer cell line was obtained from the American Tissue Culture Collection (ATCC, HTB-22^−^) and cultured in Eagle’s Minimum Essential Medium (MEM, M4655, Sigma) supplemented with 10% fetal bovine serum (FBS, 10500, Gibco, Lot 08G3057K) and 1% antibiotics (PenStrep, P4333, Sigma). Cells were maintained in a 5% CO_2_ atmosphere at 37°C. Authenticity was verified at the DNA sequencing facility of The University of Manchester by STR analysis. Mouse embryonic fibroblasts (MEFs) were extracted from E12.5 CD1 embryos following a standard protocol(39) and maintained in Dubelco’s Modified Eagles Medium (DMEM, D6429, Sigma) supplemented with 10% FBS and 1% antibiotics, in a 5% CO_2_ atmosphere at 37°C. They were used for a maximum of three passages.

### Cell exposure to GO

Cells were grown on 6-well tissue culture treated plates (3516, Corning) and, when confluency reached 70%, exposed to different subtoxic concentrations of GO (5, 10 and 50 μg/ml) in the absence of serum proteins. FBS was added 4 hours later, to reach 10% concentration, and cells were lysed 24 h after the initial exposure to interrogate gene expression. The control group was FBS-starved for the same amount of time, but was not exposed to GO. Three biological replicated were included in each group (n=3).

### MIQE guidelines

Gene expression analyses in this study adhered to the Minimum Information for Publication of Quantitative Real-Time PCR Experiments (MIQE) quidelines(21) to promote transparency and ensure the reliability of the results. All procedures were performed in the investigators’ laboratory, apart from the assessment of RNA quality with Agilent TapeStation (see below), that took place in the Genomic Core Facility at The University of Manchester. A MIQE checklist is provided in **Table S1.** Experimental details related to all steps involved in gene expression analyses are provided below.

### RNA extraction

Total RNA was isolated with the silica spin column-based PureLink^®^ RNA mini kit (12183025, Invitrogen), following the manufacturer’s instructions. Cells were lysed in 300 μl lysis buffer provided in the kit and supplemented with 1% β-mercaptoethanol (M6250, Sigma). DNAse treatment was performed on-column with Purelink DNAse set (12185010, Invitrogen). RNA was eluted in 45 μl nuclease-free water and stored at −80°C for no more than one month until further use. Samples were defrosted on ice and cDNA synthesis was performed immediately after thawing.

### Assessment of RNA integrity

RNA yield and quality were initially assessed by spectrophotometry with BioPhotometer Plus (Eppendorf). RNA concentrations and A_260/280_ and A_260/230_ ratios for each sample are reported in **Tables S2 and S3**. RNA integrity was further analyzed in Agilent 2200 TapeStation (Agilent Genomics). Briefly, samples were diluted to a final concentration within the 30-500 ng/μl range. 1 μl of the diluted RNA was denatured for 3 min at 72°C in 5 μl RNA Screen Tape Sample Buffer (5067-5577, Agilent Genomics). Samples were cooled for 2 min on ice and run in RNA Screen Tape (5067-5576, Agilent Genomics). 28S/18S ratios and RIN scores are reported in **Table S3**. Electrophoresis bands are shown in **Figure S1**.

### cDNA synthesis

1 μg RNA was converted into cDNA with the High-Capacity cDNA Reverse Transcription Kit (4368814, ThermoFisher Scientific) that includes random primers and the reverse transcriptase MultiScribe^−^. Reverse transcription (20 μl volume, 50 U reverse transcriptase) was performed in triplicate for each sample according to the following steps: 25°C for 10 min, 37°C for 120 min, 85°C for 5 min, cool down to 4°C. cDNA was stored at −20°C until qPCR was performed.

### Primer design

Details of the primers used in this study are reported in **Tables S4 and S5**. Primers were designed with Primer Basic Local Alignment Search Tool (Primer BLAST) adhering to the following requisites: amplicon size 75-200 bp, GC content 50-65%, ≤3 G or C repetitions, ≤4 base repetitions, melting temperature (Tm) 55-65°C. When gene targets had several splicing variants (including predicted variants), primer pairs were designed to amplify all of them at the same product length. Each primer pair was verified with Blast tool (NCBI) to confirm the its specificity for the desired target. Primers were synthetized by Sigma (UK) and purified by desalting.

### Real-time quantitative Polymerase Chain Reaction (qPCR)

2 ul of cDNA were used for each 20 μl qPCR reaction, which was performed with SYBR green chemistry (PowerUp^−^ SYBR^−^ Green Master Mix, A25742, Applied Biosystems) that contains Dual-lock^−^ *Taq* DNA polymerase. Reactions were loaded manually (in technical duplicates) on white-wall, clear-well, hard-shell 96-well plates (HSP9601, Biorad) and run on a CFX96 Touch real-time PCR detection system (Biorad) according to the following protocol: 2 min at 50°C, 2 min at 95°C, (15 sec at 95°C and 1 min at 60°C) x 40 cycles. The reaction was followed by a melt curve analysis, as specified by the manufacturer, to confirm amplification of a single product. No amplification was detected in non-template controls (NTC) and non-reverse transcription (NRT) controls. Cq<40 for all detections, which evidenced establishment of the limit of detection. Each gene was assessed in the same run for the totality of the samples, to avoid inter-run variability. Cq values were determined with the Single Threshold mode in the CFX Manager software (Biorad). PCR efficiencies (E) were calculated from the slope of a linear regression of the Ct values obtained from a dilution series of the starting cDNA, following the equation 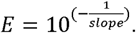

### Descriptive statistics of Cq values (Bestkeeper algorithm)

Cq values from technical duplicates were averaged and the Bestkeeper Excel tool was used to retrieve descriptive statistics (n=12). Variation was expressed as standard deviation (SD) and coefficient of variance (CV), the latter calculated as the percentage of the Cq SD to the Cq mean. Bestkeeper index for data normalization was calculated as the geometric mean of the most stable candidate genes, identified by repeated pair-wise correlation analysis, as described by Pfaffl *et al(22*)

### Assessment and ranking of reference gene stability (NormFinder algorithm)

The model-based NormFinder algorithm, as described by Andersen et al(23), was used to assess the stability of the expression of ten candidate reference genes across all experimental groups. Cq values (n=12) were transformed to relative quantities, according to the formula: E^(lowest Cq - Cq)^, that takes into account the efficiency of the PCR reaction (E) and uses the lowest Cq as a calibrator. Stability values were calculated for each candidate housekeeping gene, taking into account intra and intergroup variation. The stability value of the best combination of two genes was also calculated.

### Relative gene expression analysis

Relative gene expression was calculated following the Livak method(10). As calibrator, either the geometric mean of Cqs of two reference genes indicated by NormFinder algorithm, or the Bestkeeper index, was used. All data was normalized to the control (untreated) group. Error was propagated according to the formula: 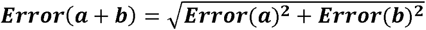

### Statistical analysis of relative gene expression data

Differences in gene expression among experimental groups were assessed with OriginPro v9.1, using ΔCq data. First, normal distribution and homogeneity of variances were confirmed (Levene’s test). Differences between groups were assessed by one-way ANOVA and Tukey’s post-hoc test. P values □0.05 were considered significant.

## Declarations

## Availability of data and materials

The datasets used and/or analyzed during the current study are available from the corresponding author on reasonable request.

## Acknowledgements

The authors would like to acknowledge the staff in the Faculty of Biology, Medicine and Health Genomic Facility for their assistance.

## Author contributions

IdL designed and performed the experiments. IdL and KK discussed the data and wrote the manuscript.

## References

1. Kostarelos K. Translating graphene and 2D materials into medicine. Nature Reviews Materials. 2016;1.

2. Bitounis D, Ali-Boucetta H, Hong BH, Min DH, Kostarelos K. Prospects and challenges of graphene in biomedical applications. Adv Mater. 2013 Apr 24;25(16):2258-68.

3. Zhao H, Ding R, Zhao X, Li Y, Qu L, Pei H, et al. Graphene-based nanomaterials for drug and/or gene delivery, bioimaging, and tissue engineering. Drug Discov Today. 2017 Sep;22(9):1302-17.

4. Bussy C, Ali-Boucetta H, Kostarelos K. Safety Considerations for Graphene: Lessons Learnt from Carbon Nanotubes. Accounts Chem Res. 2013 Mar 19;46(3):692-701.

5. Chatterjee N, Eom HJ, Choi J. A systems toxicology approach to the surface functionality control of graphene-cell interactions. Biomaterials. 2014 Jan;35(4):1109-27.

6. Li Y, Wu Q, Zhao Y, Bai Y, Chen P, Xia T, et al. Response of microRNAs to in vitro treatment with graphene oxide. ACS nano. 2014 Mar 25;8(3):2100-10.

7. Orecchioni M, Jasim DA, Pescatori M, Manetti R, Fozza C, Sgarrella F, et al. Molecular and Genomic Impact of Large and Small Lateral Dimension Graphene Oxide Sheets on Human Immune Cells from Healthy Donors. Adv Healthc Mater. 2016 Jan 21;5(2):276-87.

8. Orecchioni M, Bedognetti D, Newman L, Fuoco C, Spada F, Hendrickx W, et al. Single-cell mass cytometry and transcriptome profiling reveal the impact of graphene on human immune cells. Nat Commun. 2017 Oct 24;8(1):1109.

9. VanGuilder HD, Vrana KE, Freeman WM. Twenty-five years of quantitative PCR for gene expression analysis. Biotechniques. 2008 Apr;44(5):619-26.

10. Livak KJ, Schmittgen TD. Analysis of relative gene expression data using real-time quantitative PCR and the 2(-Delta Delta C(T)) Method. Methods. 2001 Dec;25(4):402-8.

11. Pfaffl MW. A new mathematical model for relative quantification in real-time RT-PCR. Nucleic Acids Res. 2001 May 1;29(9):e45.

12. Huggett J, Dheda K, Bustin S, Zumla A. Real-time RT-PCR normalisation; strategies and considerations. Genes Immun. 2005 Jun;6(4):279-84.

13. Kozera B, Rapacz M. Reference genes in real-time PCR. J Appl Genet. 2013 Nov;54(4):391-406.

14. Ali H, Du Z, Li X, Yang Q, Zhang YC, Wu M, et al. Identification of suitable reference genes for gene expression studies using quantitative polymerase chain reaction in lung cancer in vitro. Mol Med Rep. 2015 May;11(5):3767-73.

15. Lemma S, Avnet S, Salerno M, Chano T, Baldini N. Identification and Validation of Housekeeping Genes for Gene Expression Analysis of Cancer Stem Cells. PLoS One. 2016;11(2):e0149481.

16. Glare EM, Divjak M, Bailey MJ, Walters EH. beta-Actin and GAPDH housekeeping gene expression in asthmatic airways is variable and not suitable for normalising mRNA levels. Thorax. 2002 Sep;57(9):765-70.

17. Waxman S, Wurmbach E. De-regulation of common housekeeping genes in hepatocellular carcinoma. BMC Genomics. 2007 Jul 18;8:243.

18. Wierschke S, Gigout S, Horn P, Lehmann TN, Dehnicke C, Brauer AU, et al. Evaluating reference genes to normalize gene expression in human epileptogenic brain tissues. Biochem Biophys Res Commun. 2010 Dec 17;403(3-4):385-90.

19. Gubern C, Hurtado O, Rodriguez R, Morales JR, Romera VG, Moro MA, et al. Validation of housekeeping genes for quantitative real-time PCR in in-vivo and in-vitro models of cerebral ischaemia. BMC Mol Biol. 2009 Jun 16;10:57.

20. Everaert BR, Boulet GA, Timmermans JP, Vrints CJ. Importance of suitable reference gene selection for quantitative real-time PCR: special reference to mouse myocardial infarction studies. PLoS One. 2011;6(8):e23793.

21. Bustin SA, Benes V, Garson JA, Hellemans J, Huggett J, Kubista M, et al. The MIQE guidelines: minimum information for publication of quantitative real-time PCR experiments. Clin Chem. 2009 Apr;55(4):611-22.

22. Pfaffl MW, Tichopad A, Prgomet C, Neuvians TP. Determination of stable housekeeping genes, differentially regulated target genes and sample integrity: BestKeeper‐‐Excel-based tool using pair-wise correlations. Biotechnol Lett. 2004 Mar;26(6):509-15.

23. Andersen CL, Jensen JL, Orntoft TF. Normalization of real-time quantitative reverse transcription-PCR data: a model-based variance estimation approach to identify genes suited for normalization, applied to bladder and colon cancer data sets. Cancer research. 2004 Aug 1;64(15):5245-50.

24. Vandesompele J, De Preter K, Pattyn F, Poppe B, Van Roy N, De Paepe A, et al. Accurate normalization of real-time quantitative RT-PCR data by geometric averaging of multiple internal control genes. Genome Biol. 2002 Jun 18;3(7):RESEARCH0034.1-RESEARCH.11.

25. Vincent M, de Lazaro I, Kostarelos K. Graphene materials as 2D non-viral gene transfer vector platforms. Gene Ther. 2017 Jan 05.

26. de Jonge HJ, Fehrmann RS, de Bont ES, Hofstra RM, Gerbens F, Kamps WA, et al. Evidence based selection of housekeeping genes. PLoS One. 2007 Sep 19;2(9):e898.

27. Ali-Boucetta H, Bitounis D, Raveendran-Nair R, Servant A, Van den Bossche J, Kostarelos K. Purified graphene oxide dispersions lack in vitro cytotoxicity and in vivo pathogenicity. Adv Healthc Mater. 2013 Mar;2(3):433-41.

28. Mukherjee SP, Lozano N, Kucki M, Del Rio-Castillo AE, Newman L, Vazquez E, et al. Detection of Endotoxin Contamination of Graphene Based Materials Using the TNF-alpha Expression Test and Guidelines for Endotoxin-Free Graphene Oxide Production. Plos One. 2016 Nov 23;11(11).

29. Mukherjee SP, Lazzaretto B, Hultenby K, Newman L, Rodrigues AF, Lozano N, et al. Probing graphene oxide-induced plasma membrane perturbations in neutrophils using time-of-flight secondary ion mass spectroscopy. Chem 2017;accepted.

30. Dheda K, Huggett JF, Bustin SA, Johnson MA, Rook G, Zumla A. Validation of housekeeping genes for normalizing RNA expression in real-time PCR. Biotechniques. 2004 Jul;37(1):112-4, 6, 8-9.

31. Barber RD, Harmer DW, Coleman RA, Clark BJ. GAPDH as a housekeeping gene: analysis of GAPDH mRNA expression in a panel of 72 human tissues. Physiol Genomics. 2005 May 11;21(3):389-95.

32. Feng LZ, Yang XZ, Shi XZ, Tan XF, Peng R, Wang J, et al. Polyethylene Glycol and Polyethylenimine Dual-Functionalized Nano-Graphene Oxide for Photothermally Enhanced Gene Delivery. Small. 2013 Jun 10;9(11):1989-97.

33. Zhi F, Dong HF, Jia XF, Guo WJ, Lu HT, Yang YL, et al. Functionalized Graphene Oxide Mediated Adriamycin Delivery and miR-21 Gene Silencing to Overcome Tumor Multidrug Resistance In Vitro. Plos One. 2013 Mar 20;8(3).

34. Huang YP, Hung CM, Hsu YC, Zhong CY, Wang WR, Chang CC, et al. Suppression of Breast Cancer Cell Migration by Small Interfering RNA Delivered by Polyethylenimine-Functionalized Graphene Oxide. Nanoscale Res Lett. 2016 Dec;11(1):247.

35. Choi HY, Lee TJ, Yang GM, Oh J, Won J, Han J, et al. Efficient mRNA delivery with graphene oxide-polyethylenimine for generation of footprint-free human induced pluripotent stem cells. J Control Release. 2016 Aug 10;235:222-35.

36. Zhang L, Zhou Q, Song W, Wu K, Zhang Y, Zhao Y. Dual-Functionalized Graphene Oxide Based siRNA Delivery System for Implant Surface Biomodification with Enhanced Osteogenesis. ACS Appl Mater Interfaces. 2017 Oct 11;9(40):34722-35.

37. Yin F, Hu K, Chen Y, Yu M, Wang D, Wang Q, et al. SiRNA Delivery with PEGylated Graphene Oxide Nanosheets for Combined Photothermal and Genetherapy for Pancreatic Cancer. Theranostics. 2017;7(5):1133-48.

38. De Spiegelaere W, Dern-Wieloch J, Weigel R, Schumacher V, Schorle H, Nettersheim D, et al. Reference gene validation for RT-qPCR, a note on different available software packages. PLoS One. 2015;10(3):e0122515.

39. Michalska AE. Isolation and propagation of mouse embryonic fibroblasts and preparation of mouse embryonic feeder layer cells. Curr Protoc Stem Cell Biol. 2007 Oct;3:1C.3.1-C.3.17.

